# To mate, or not to mate: The evolution of reproductive diapause facilitates insect radiation into African savannahs in the Late Miocene

**DOI:** 10.1101/693812

**Authors:** Sridhar Halali, Paul M. Brakefield, Steve C. Collins, Oskar Brattström

## Abstract

1. Many tropical environments experience cyclical seasonal changes, frequently with pronounced wet and dry seasons, leading to a highly uneven temporal distribution of resources. Short-lived animals inhabiting such environments often show season-specific adaptations to cope with alternating selection pressures.
2. African *Bicyclus* butterflies show strong seasonal polyphenism in a suite of phenotypic and life-history traits, and their adults are thought to undergo reproductive diapause associated with the lack of available larval host plants during the dry season.
3. Using three years of longitudinal field data for three species in Malawi, dissections demonstrated that one forest species reproduces continuously whereas two savannah species undergo reproductive diapause in the dry season, either with or without pre-diapause mating. Using additional data from field-collected and museum samples, we then documented the same three mating strategies for a further 37 species.
4. Phylogenetic analyses indicated that the ancestral state was a non-diapausing forest species, and that habitat preference and mating strategy evolved in a correlated fashion.
5. *Bicyclus* butterflies underwent rapid diversification during the Late Miocene, coinciding with expansions into more open savannah habitat. We conclude that the ability to undergo reproductive diapause was a key trait that facilitated colonization and eventual radiation into savannahs in the Late Miocene.

## 1. Introduction

Many tropical environments experience cyclical seasonal changes. Alternating wet and dry seasons are especially widespread, the former with abundant resources (e.g., host plants for phytophagous insects), whereas the latter are characterised by minimal resources (Denlinger, 1986; Jones, 1987; Tauber, Tauber, & Masaki, 1986). Thus, animals inhabiting such environments face dramatic changes in selection pressures with selection typically favouring traits that increase reproduction in the wet season, and survival in the dry season. Insects in seasonal environments often synchronise their reproduction with the phenology of associated plants to optimise the abundance and quality of resources available both for larvae and adults (Denlinger, 1986; Tauber et al., 1986).

Many tropical insects enter diapause at a particular life stage (egg, larval, pupal, or adult) or move to more favourable habitats to escape extreme environments of the dry season (Denlinger, 1986; Tauber et al., 1986). Diapause is an arrest of development or ontogeny coupled with co-ordinated changes involving numerous physiological and life history traits such as low metabolic activity, increased fat deposits and longevity, and altered or reduced behavioral acitivity (Hahn & Denlinger, 2007, 2010; Tauber et al., 1986; Tatar & Yin, 2001). Together these traits are considered to facilitate survival in the resource-depleted environment (Tauber et al., 1986). Although diapause can be a highly effective strategy allowing temporal escape of unfavourable conditons, it is likely to come with costs including a higher mortality and a significant reduction in post-diapause fecundity (Ellers & van Alphen, 2002; Hahn & Denlinger, 2007). From a macroevolutionary perspective, strategies such as diapause can be a key life-history trait which can enable colonization of environments where habitat quality fluctuates temporally (Furness, Reznick, Springer, & Meredith, 2015; Panov & Caceres, 2004).

Mycalesina butterflies (Nymphalidae: Satyrinae: Satyrini) are distributed throughout the Old-World tropics and have undergone parallel radiations in Southeast Asia, Sub-Saharan mainland Africa, and Madagascar (Aduse-Poku et al., 2015; Kodandaramaiah et al., 2010). The adults of many Mycalesina butterflies show seasonal polyphenism in wing pattern with alternative wet and dry season forms produced through developmental phenotypic plasticity (Brakefield & Larsen, 1984; Brakefield & Reitsma, 1991). The wet season form has characteristic conspicuous eyespots with concentric coloured rings positioned along the wing margins. These have been shown to function in deflecting predator attacks to the wing margins and away from more vulnerable body parts (Lyytinen, Brakefield, & Mappes, 2003; Lyytinen, Brakefield, Lindström, & Mappes, 2004; Olofsson, Vallin, Jakobsson, & Wiklund, 2010; Prudic, Stoehr, Wasik, & Monteiro, 2015). In contrast, the dry season form generally lack these eyespots, and they rely mainly on crypsis when at rest on dead leaf litter for survival against predators hunting by sight (Lyytinen et al., 2003, 2004; Prudic et al., 2015). Along with the wing pattern changes, both forms differ in a suite of other morphological, behavioural, physiological, and life-history traits, as an integrated response which is linked by a common hormonal switch as has been extensively explored in laboratory settings in a single species *Bicyclus anynana* (Oostra et al., 2010, 2014; van Bergen & Beldade, 2019; van Bergen et al., 2017). Such an integrated system helps to maintain a season-specific adaptive phenotype by making use of environmental cues to predict approaching seasonal shifts (Brakefield & Reitsma, 1991; Kooi & Brakefield, 1999).

Species of the African Mycalesina genus *Bicyclus* inhabit both forests and open savannah habitats with forest species probably reflecting the ancestral state (van Bergen, 2015). They mainly use grasses (Poaceae) as their larval host plants with a few records of utilising plants belonging to Zingiberaceae and Marantaceae family (Larsen, 2005). Stable isotope analysis has also shown that forest and savannah species of *Bicyclus* prefer C_3_ or C_4_ grasses, respectively, as larval host plants (van Bergen et al., 2016). C_3_ grasses are usually forest-linked, while C_4_ grasses predominate in savannah grasslands, the former being the ancestral state (Edwards et al., 2010). Savannah grasslands began to open up from forest-dominated habitats in Africa during the Late Miocene and Pliocene (Cerling et al., 1997; Edwards et al., 2010; Jacobs, 2004; Osborne, 2008). Interestingly, *Bicyclus* (and the Satyrini tribe in general) showed a burst of radiation at this time suggesting that expansion of savannahs opened up new niches and subsequent colonization of these habitats may have resulted in the rapid radiation of these butterflies (Aduse-Poku et al. 2015; Brakefield, 2010; Kodandaramaiah et al., 2010; Peña & Wahlberg, 2008). However, colonizing highly seasonal savannahs from more stable forest habitats may only have been possible through the evolution of key adaptations (Brakefield, 2010).

One such potential key trait, that has been hypothesised to be used by *Bicyclus*, is the capacity of adults to undergo reproductive diapause during the dry season. During reproductive diapause the development of oogenesis, vitellogenesis, accessory glands, and mating behaviour is arrested (Tauber et al., 1986; Tatar & Yin, 2001). In *Bicyclus* butterflies, *B. anynana* in particular, when entering reproductive diapause, females arrest reproduction coupled with changes in other traits such as cryptic colour pattern, large body size, increased fat deposits, increased longevity, and reduced behavioural activity for the entire dry season until the onset of the rains (Brakefield & Reitsma, 1991; Oostra et al., 2014; van Bergen et al. 2017). This would enable a suspension of reproductive effort, while still allowing opportunistic feeding and dispersal behaviour by adults and allowing a rapid switch to reproduction as the rains arrive and larval host plants become available again. However, this hypothesis has rarely been tested in the field. A survey on three Australian Mycalesina butterflies showed that species exhibit different mating strategies in the dry season in a habitat-dependent manner. Thus, the savannah species undergo reproductive diapause, while forest species reproduce continuously in line with a constant availability of host plants throughout the year (Braby, 1995; Moore, 1986). Another study in Zomba, Malawi, examined 49 females of *Bicyclus safitza* from the peak of the dry season and found that 36% of these carried spermatophores, a clear evidence of having mated. They also had mature eggs in the dry season, apparently indicating that not all adults of this species undergo reproductive diapause (Brakefield & Reitsma, 1991).

In this study we explore these patterns further. Firstly, we use three years of continuously collected field samples from Zomba, Malawi, and document three different mating strategies that females use in the dry season: no reproductive diapause or reproductive diapause, either with mating before diapause (pre-diapause mating) or with mating only after diapause (complete reproductive diapause). Secondly, by examining museum samples for an additional 37 species we demonstrate similar three mating strategies across all the species. Further, phylogenetic analyses show that habitat preference and mating strategy evolve in a correlated fashion with non-dipausing forest species as the ancestral state. We also construct an evolutionary pathway which suggests that the gain of the ability to show reproductive diapause is a key trait in facilitating colonization of savannahs in these butterflies in the Late Miocene.

## 2. Materials and methods

### 2.1 Using long-term field samples to investigate mating strategies in the dry season

We used field-collected samples to explore the potential mating and reproductive strategies used by *Bicyclus* butterflies in the dry season. Butterflies were collected daily using three fruit-baited traps (fermented banana) operated over three years from the beginning of the dry season in 1995 until the end of the wet season in 1998 in Zomba, Malawi, and all the collected samples were dried after collection and stored in glassine envelopes. These samples have previously been used in another study (van Bergen et al., 2016) where the specimens we now refer to as *B. campina* were treated as *B. vansoni*. Our reassignment of their species identity was based on the geographic location and the wing morphology, as well as some doubts about the validity of the species *B. vansoni* (Oskar Brattström, personal communication).

Three species, *B. campina, B. safitza*, and *B. ena*, were abundant in the collections and well represented across all the three years of sampling. *B. campina* mainly prefers forest habitats, including forest edges (Brakefield & Reitsma, 1991; van Bergen et al., 2016; Windig et al., 1994). *B. safitza* is one of the most abundant and widely distributed species of *Bicyclus* which mainly occurs in open savannah and degraded forests (Brakefield & Reitsma, 1991; van Bergen et al., 2016; Windig, Brakefield, Reitsma, & Wilson, 1994). *B. ena* mainly occurs in the open savannahs with a preference for dry rocky habitats (Brakefield & Reitsma, 1991; van Bergen et al., 2016; Windig et al., 1994).

In butterflies, the male transfers sperm along with some nutrients in a spermatophore which is stored in the *bursa copulatrix* of the female (Burns, 1968). The presence/absence and the number of spermatophores has been widely used as a good proxy to infer mating status and mating rate from field caught samples (Burns, 1968; Braby, 1995; Larsdotter Mellström & Wiklund, 2010). Although spermatophores can become degraded upon utilization by the female, some remnants almost always remain making it possible to accurately count them (Burns, 1968; but see Walters, Stafford, Hardcastle, & Jiggins, 2012).

We aimed to dissect at least 10 females of each species per month throughout the whole sample period, but enough individuals were not always available in December to January. For carrying out dissections, we first soaked the abdomens in 0.4 ml of 5% Trisodium Phosphate and Ilfotol solution (master mix of 50 ml containing 48 ml 5% of Trisodium Phosphate in water and 2 ml Ilfotol, modified from Burns (1968)), and kept them at room temperature overnight. During the dissections we counted spermatophores and noted the presence/absence of mature (yolk filled) eggs.

To classify butterflies into seasonal forms, we photographed all the individuals and measured the relative area of the CuA1 eyespot and hindwing wing area using a custom written macro in the ImageJ 1.8.0 software (Schneider, Rasband, & Eliceiri, 2012) (supplementary material, Figure S1). We plotted histograms to explore the distribution of eyespot areas, and butterflies were then classified as the wet or dry form by setting a threshold (see van Bergen et al., 2016, supplementary material, Figure S2). We estimated repeatabilities by re-measuring a sub-sample of 60 butterflies on different dates giving R^2^ values of 0.98 and 0.96 for eyespot area and wing area, respectively.

### 2.2 Sampling populations across habitats to test variation in the mating strategy

Insect mating strategies can change across populations inhabiting different habitats (Dingle, 1968; Denlinger, 1986; Tauber et al., 1986). To test whether this happens in *Bicyclus*, we used samples collected from a field survey in Ghana conducted in the early dry season from 5^th^ to 19^th^ December 2018. We sampled two selected species, *B. funebris* and *B. vulgaris*, from multiple populations in Ghana across different habitats ranging from wet rainforest to dry forest edges. Similarly, we checked three populations of *B. anynana* from personal collections and museum samples from South Africa, Tanzania and Zimbabwe to test whether there is geographic variation in mating strategy. Samples from South Africa and Zimbabwe were sampled from a single dry season but from different years, while samples from Tanzania were combined across different years.

### 2.3 Sampling of species across the phylogeny

In addition to the three *Bicyclus* species from Zomba, we surveyed smaller numbers of samples of 37 additional *Bicyclus* species collected in the middle of the dry season. We primarily used material from the collections of the African Butterfly Research Institute (ABRI) in Nairobi, Kenya, and from field surveys across several sites in Ghana (December 2018) and Kakamega forest in Kenya (February 2019), and from the personal collection of Oskar Brattström. We aimed to sample ten specimens per species from museum samples and up to 25 samples per site in the field. For sampling the museum specimens from the middle of the dry season, we first checked annual climate graphs for the specific locality to identify a suitable time period as distant as possible from any substantial rainfall. Analysis of the long-term samples from Zomba suggests that a small number of females from the peak dry season (July to October) is sufficient to identify the mating strategy (see Results) and all our additional species could be fitted into the same classifications. For some of the additional species it was not possible to analyse only individuals from a single location, and therefore we occasionally combined samples from multiple sites and sometimes countries. This did not affect assigning species to specific mating strategies as populations exhibited the same strategy across populations (see Results).

We sampled a total of 228 specimens from 23 species from the collections at ABRI (mean per species = 9.9; range = 6-19) (supplementary material, Table S1). The field sampling in Ghana provided 334 specimens from 11 species (mean = 30.7; range = 7-100) (supplementary material, table S2) and the Kenyan sampling provided 73 specimens from 5 species (mean = 14.6; range = 7-37) (supplementary material, Table S1). We also included 25 Nigerian and South-African samples from personal collections of two species (mean = 12.5; range = 9-16) (supplementary material, table S1). Combining these data with the three species from our long-term collection, our final dataset consisted of 40 species of *Bicyclus*.

### 2.4 Classification of mating and reproductive strategies, and habitat preferences

Following examination of the total data set for 40 species, we used the percentage of females with presence of spermatophores and mature eggs to categorize species into one of three dry season mating and reproductive strategies: (1) non-diapausing with continuous reproduction (henceforth non-diapausing) where females have spermatophores and mature eggs in the dry season, (2) completely diapausing with no spermatophores or eggs in the middle of the dry season and mating occurs only at the end (in a few cases a small proportion of mid-season females have spermatophores) (3) pre-diapause mating where females mate in the early dry season but do not develop eggs until the end of the dry season (see Kato, 1986). In this strategy females may mate again in the late dry season (Kato, 1986).

For categorising species into one of the three mating strategies we considered a threshold of 50%. That is, if ≥50% of females had spermatophores and mature eggs then the species was classified as non-diapausing. Similarly, if < 50% of females had spermatophores and eggs it was considered as a diapausing species. In cases where ≥50% of females had spermatophores but less than half of those had eggs it was considered as a pre-diapause mating species. Most of the species exhibited clear mating strategies except for two forest species, *B. dorothea* and *B. procora*, which had 50% of specimens containing spermatophores and eggs. There is evidence that time to reproductive maturity is longer in forest than open habitat species and hence it is possible that these species may take a longer time to mate and develop eggs (Braby, 2002). Based on this reasoning we classified both these species as non-diapausing.

In the case of one species *B. larseni*, one of the smallest of all *Bicyclus* species, it was difficult to find spermatophores in 78% of specimens, but the majority had mature eggs. We classified this species as non-diapausing. This is justified by the observation that it is costly for the female to have mature eggs without mating in the dry season, as heavy abdomens can affect flight and hence increase predation (Marden & Chai, 1991).

Habitat preference was classified for each species as one of three categories: forest-restricted, forest-dependent, or savannah. This was based on literature (Condamin, 1973; Kielland, 1990; Larsen, 2005; van Bergen, 2015), and our own extensive experience from field-work. Forest-dependent species usually occur in forest fringes and can tolerate some degree of habitat degradation but are generally not found in true savannah habitats. While, forest-restricted species are generally found in rainforests.

### 2.5 Ancestral state reconstruction

We used an updated recent ultrametric phylogeny of Mycalesina (O. Brattström, K. Aduse-Poku, E. van Bergen, V. French, & P.M. Brakefield, *under review*) which is built on previous phylogenies (Aduse-Poku et al., 2015, Aduse-Poku, Brakefield, Wahlberg, & Brattström, 2017) for our phylogenetic analyses after trimming it down to those 40 species surveyed here. We performed ancestral state reconstruction to visualise the evolution of mating strategies and habitat preference applying Bayesian stochastic mapping of ancestral states (Bollback, 2006; Huelsenbeck, Nielsen, & Bollback, 2003) by simulating characters 1000 times on maximum clade credibility tree using *make.simmap* function implemented in Phytools Ver.0.6.60 (Revell, 2012). For both investigated traits we fitted three models of character evolution: equal rates, symmetric, and all rates different model. The models were fitted using the *fitMk* function in Phytools and the best fitting model was chosen based on the Akaike Information Criteria (AIC) weights and was then used for the ancestral state reconstruction. Unless otherwise stated, we performed all the analyses in RStudio Ver.3.4.4. “Kite-Eating Tree” (R Core Team, 2017).

### 2.6 Testing for correlated evolution of habitat preference and mating strategies

We used DISCRETE model in BayesTraits V.3.0.1 (Meade & Pagel, 2016) to test for evidence of correlated evolution between habitat preference and mating strategy which implements Reversible Jump Markov Chain Monte Carlo (RJ MCMC) to model all the possible forward and backward transitions but does not allow simultaneous transitions in both the states (Pagel, 1994; Pagel & Meade, 2006). For the analysis of correlated evolution of discrete traits this is currently the only available method and the DISCRETE model is limited to binary traits. While, both the habitat preference and mating strategy data had three states. Therefore, we merged the diapausing and pre-diapause mating strategies into a single category. This is biologically justified because in both strategies females have immature oocytes and do not undergo active reproduction even if they mate and store spermatophores, such as in pre-diapause mating (see Kato, 1986, 1989). Similarly, for habitat preference, the forest-dependent state was merged with forest-restricted to form a single forest linked state. Forest-dependent species usually occur in the forest fringes or edges but never in true savannah habitats suggesting that forest cover is essential. Hence, merging this category with forest-restricted species is more appropriate than into savannahs.

The test for correlated evolution is performed by evaluating support for independent and dependent model. In the dependent model the rate of change of one trait depends on the other trait, while the independent model considers no such association. We ran RJ MCMC chains for ten million iterations with a burnin period of one million after which the chain was sampled every 500^th^ iteration. The chains were visually inspected for convergence. We used exponential hyperprior 0-10 and placed 1000 stepping stones each iterating 10000 times to obtain the marginal likelihood value for both the independent and dependent models. We ran the analysis three times to check for the stability of the likelihood values. We then calculated Bayes Factor which is a measure of the strength of correlated evolution using the marginal likelihood value for both models. A value between 2-6 is considered as positive evidence, while the value of 6-10 is considered strong evidence for correlated evolution (Currie & Meade, 2014).

Furthermore, using the values of transition parameters resulting from RJ MCMC, we calculated Z values for the transitions which is the proportion of times a certain transition parameter is set to zero among all sampled models (Pagel & Meade, 2006). A high proportion of zeros, and hence a high Z value, suggests that the transition between states is unlikely. Using the Z values, we constructed a pathway for habitat-dependent evolution of mating strategies.

### 2.7 Validation of co-evolutionary models

We used five approaches to validate our co-evolutionary models. (1) We tested whether our analysis is sensitive to the choice of prior by using two more priors, uniform 0-100 and exponential hyperprior 0-100. (2) We tested the effect of using different thresholds for classification of mating strategies by using an additional threshold of 10% and 20%. That is, species having spermatophores and eggs greater and lower than the threshold were classified as non-diapausing and diapausing species, respectively. And, pre-diapause mating if presence of spermatophores were greater and eggs less than the threshold. (3) Two species, *B. dorothea* and *B. procora*, in which females had 50% of spermatophores and eggs, were re-classified as diapausing species (contrary to non-diapausing in the original data-set) to test if our analysis is sensitive to this classification. (4) We tested if merging forest-dependent state with savannahs affects our conclusion regarding correlated evolution compared to the original data set where forest-dependent state was merged with forest-restricted. (5) We also tested the effect of sample size on the co-evolutionary models by randomly removing 30% of species from the data set and repeated this with ten different subsets (for similar analysis see Watts, Sheehan, Atkinson, Bulbulia, & Gray, 2016).

## 3. Results

### 3.1 Seasonality and mating patterns in the long-term data

A total of 232 *B. campina*, 373 *B. safitza*, and 206 *B. ena* females from Zomba, Malawi, were dissected. Over the whole series of samples from the peak dry season (July -October) over three annual cycles, 61% of *B. campina* females had spermatophores and eggs, while in *B. safitza* 22% and 20% of females had spermatophore and eggs, respectively. Interestingly, in *B. ena*, 88% of the females had spermatophores, but only 28% of all females had eggs (Figure 1). Reproductive activity appeared to increase at the end of the dry season just before the onset of the rain in November (Figure 1). Furthermore, in all the three species there was no significant yearly variation in the proportion of females that had spermatophores and eggs in the middle of the dry season (χ^2^ test, P>0.05). Combining all seasons for all three species, the presence of mature eggs was correlated with the presence of spermatophores in 98% of the females, but not vice-versa (i.e. spermatophores did not predict egg presence). Based on the percentage of females having spermatophores and eggs in the dry season, *B. campina* was classified as a non-diapausing species. In *B. ena*, females had spermatophores but generally lacked mature eggs, and so it was classified as a pre-diapause mating species. In *B. safitza*, only about a quarter of the females were mated and had eggs, and it was classified as a diapausing species.

**Figure 1:**
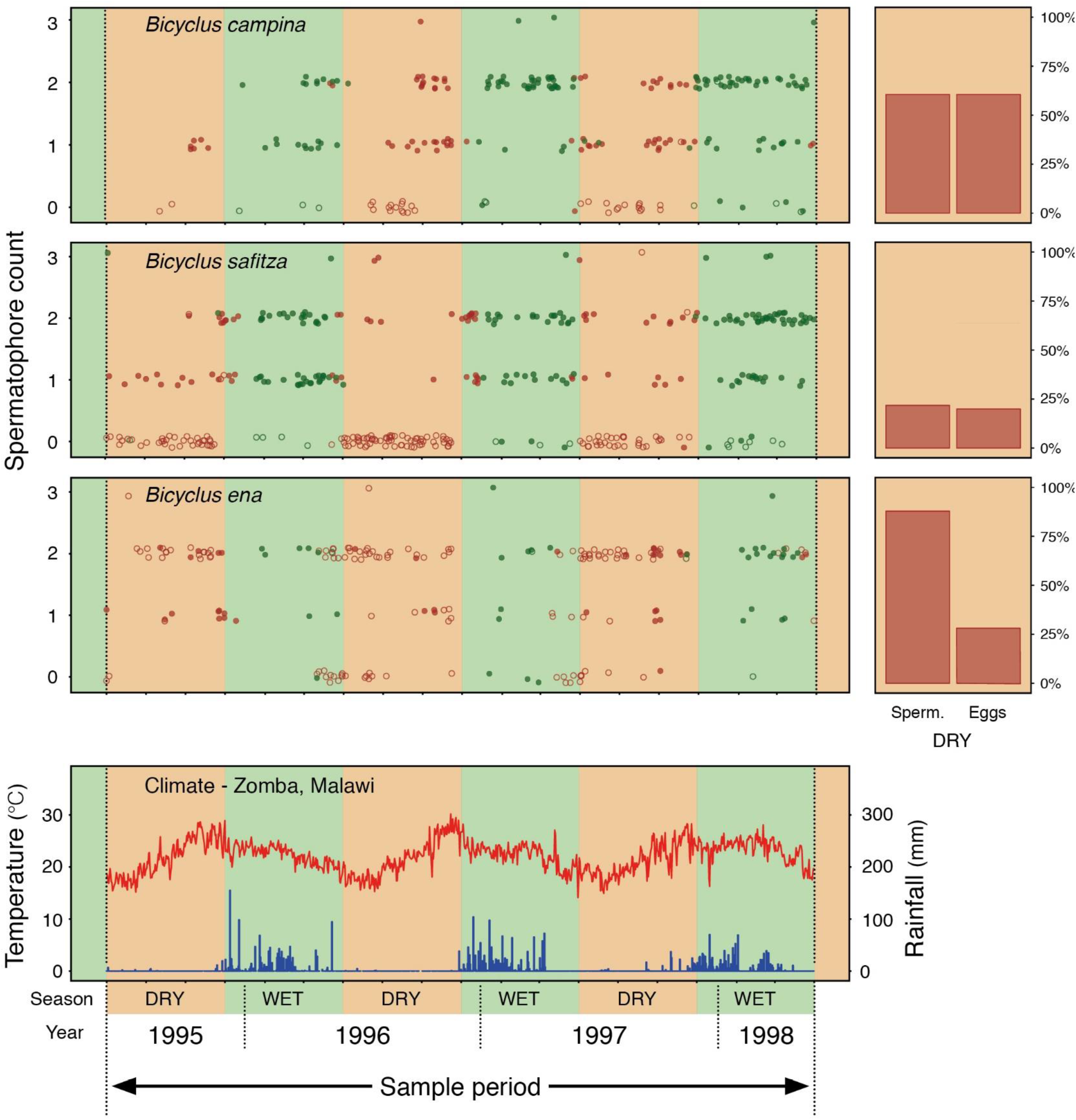
Mating strategies of females from three *Bicyclus* species, *B. safitza, B. campina* and *B. ena* across alternating wet and dry seasons in Zomba, Malawi, from July 1995–May 1998. Circles in the top three left panels represent individual butterflies (brown, dry season form; green, wet season form; open circles, mature eggs absent; filled circles, mature eggs present) with variation in temperature (red line) and rainfall (blue bars) shown in the lower panel. The seasons are marked with brown (dry) and green (wet) overlay across all panels. Note that the data points of top three left panels are jittered for clarity. The dry season is characterised by lower temperature and usually lasts from June to mid-November. In contrast, the wet season is characterised by a higher temperature and abundant rain and usually lasts from late November until late May. Stacked plots besides each species depict the percentage of females carrying spermatophores and eggs in the peak dry season (July to October).

### 3.2 Mating strategies across populations

In the two species, *B. funebris* and *B. vulgaris*, with samples from five and four localities respectively, the mating strategy (diapausing) was consistent across all populations (table 1). In addition, *B. anynana* from South Africa, Tanzania, and Zimbabwe exhibited the same mating strategy (pre-diapause mating) (table 2).

**Table 1:** Populations of two species of *Bicyclus* sampled across different habitats in Ghana. In both the species, the presence of spermatophores was always associated with the presence of mature eggs, and hence only the percentage of females having spermatophores is provided in the table (Amedzofe, slightly degraded forest; Atewa, forest; Bonkro, forest; Oporo, forest-savannah ecotone; Bobiri, forest)

|  | Amedzofe |  | Atewa |  | Bonkro |  | Bobiri |  | Oporo |  |
| --- | --- | --- | --- | --- | --- | --- | --- | --- | --- | --- |
| Species | N | % mated | N | % mated | N | % mated | N | % mated | N | % mated |
| *B. funebris* | 13 | 0 | 26 | 0 | 15 | 0 | 19 | 0 | 21 | 9.52 |
| *B. vulgaris* | 13 | 0 | 20 | 5 | - | - | 15 | 0 | 20 | 5 |

**Table 2:** Mating strategies across three populations of *B. anynana*, a species which exhibits the pre-diapause mating strategy.

|  | N | % mated | % presence<br>of eggs |
| --- | --- | --- | --- |
| South Africa | 16 | 100 | 0 |
| Tanzania | 9 | 77.77 | 22.22 |
| Zimbabwe | 10 | 100 | 0 |

### 3.3 Ancestral state reconstruction

For ancestral state reconstruction, the equal rates model showed the highest AIC weight for both habitat preferences and mating strategies (supplementary material, Table S2). Ancestral state reconstruction of mating strategies suggested that non-diapausing was the ancestral state. Similarly, for habitat preference, forest-restricted was indicated as the ancestral state (Figure 2).

**Figure 2:**
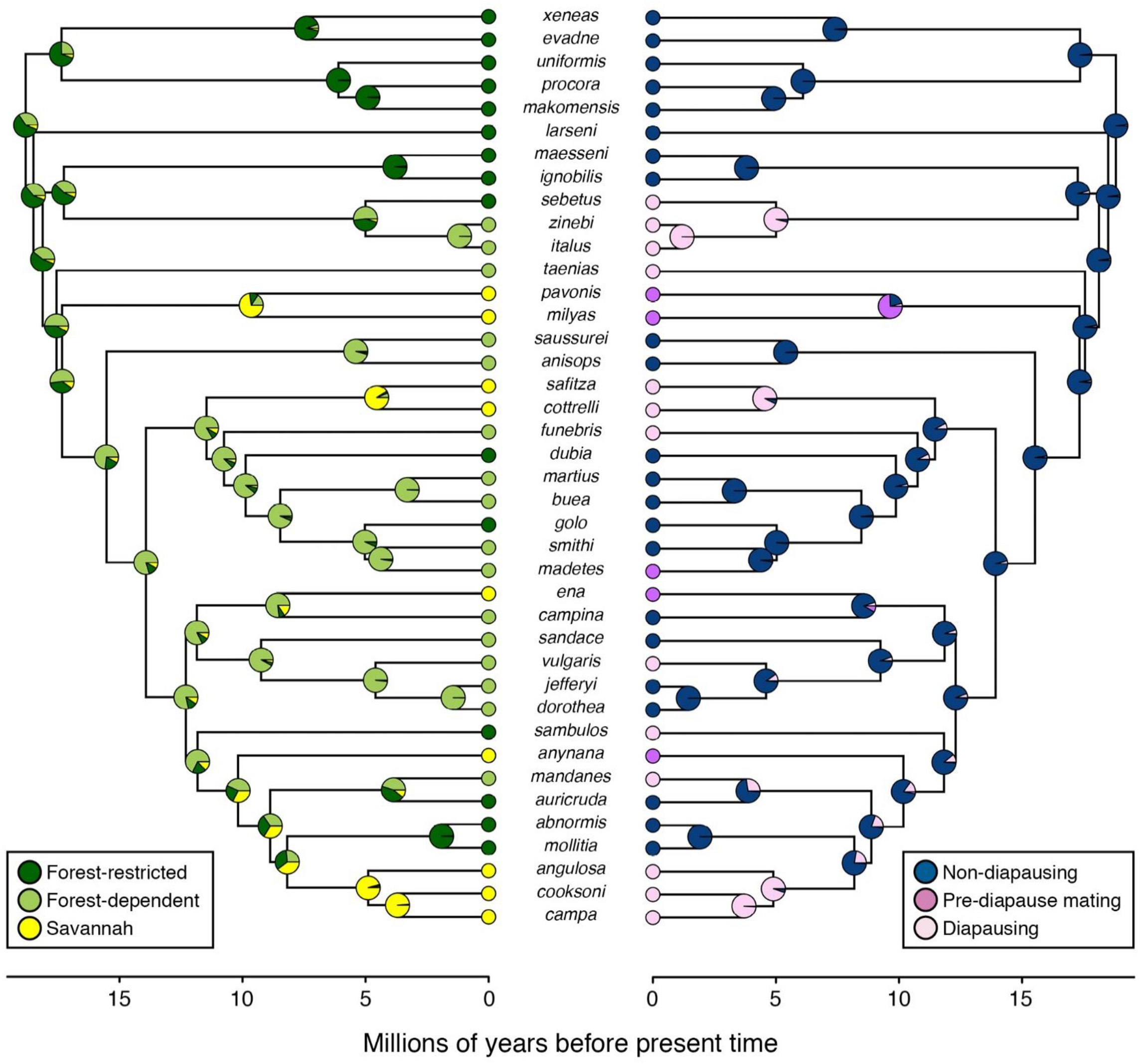
Ancestral state reconstruction for habitat preferences (left) and mating strategies (right) for species of *Bicyclus* butterfly.

### 3.4 Correlated evolution and the evolutionary pathway

Using exponential hyperprior 0-10, the dependent model of evolution provided a better fit (average marginal likelihood: −40.48) than the independent model (average marginal likelihood: −47.49), and the values were similar for three independent runs (supplementary material, table S3). The marginal likelihood value for both the models resulted in an average Bayes Factor of 14.01. The likelihood values and Bayes factor were similar for the other two priors, uniform 0-100 and exponential hyperprior 0-100, albeit slightly lower for the former prior (supplementary material, Table S3). Furthermore, the analyses conducted to test the sensitivity of co-evolutionary models to classification of habitat preference, mating strategies, and sample size, all provided consistent positive support for correlated evolution (supplementary material, Table S4–7).

The evolutionary pathway (Figure 3) shows that the transition from the ancestral state (Forest, Non-diapausing) to the derived state (Savannah, Diapausing) was generally reached through one of the two possible intermediate states (Forest, Diapausing). The transition rates leading up to the other alternative intermediate state (Savannah, Non-diapausing) were assigned to zero on most occasions and none of our sampled species show this combination of traits.

**Figure 3:**
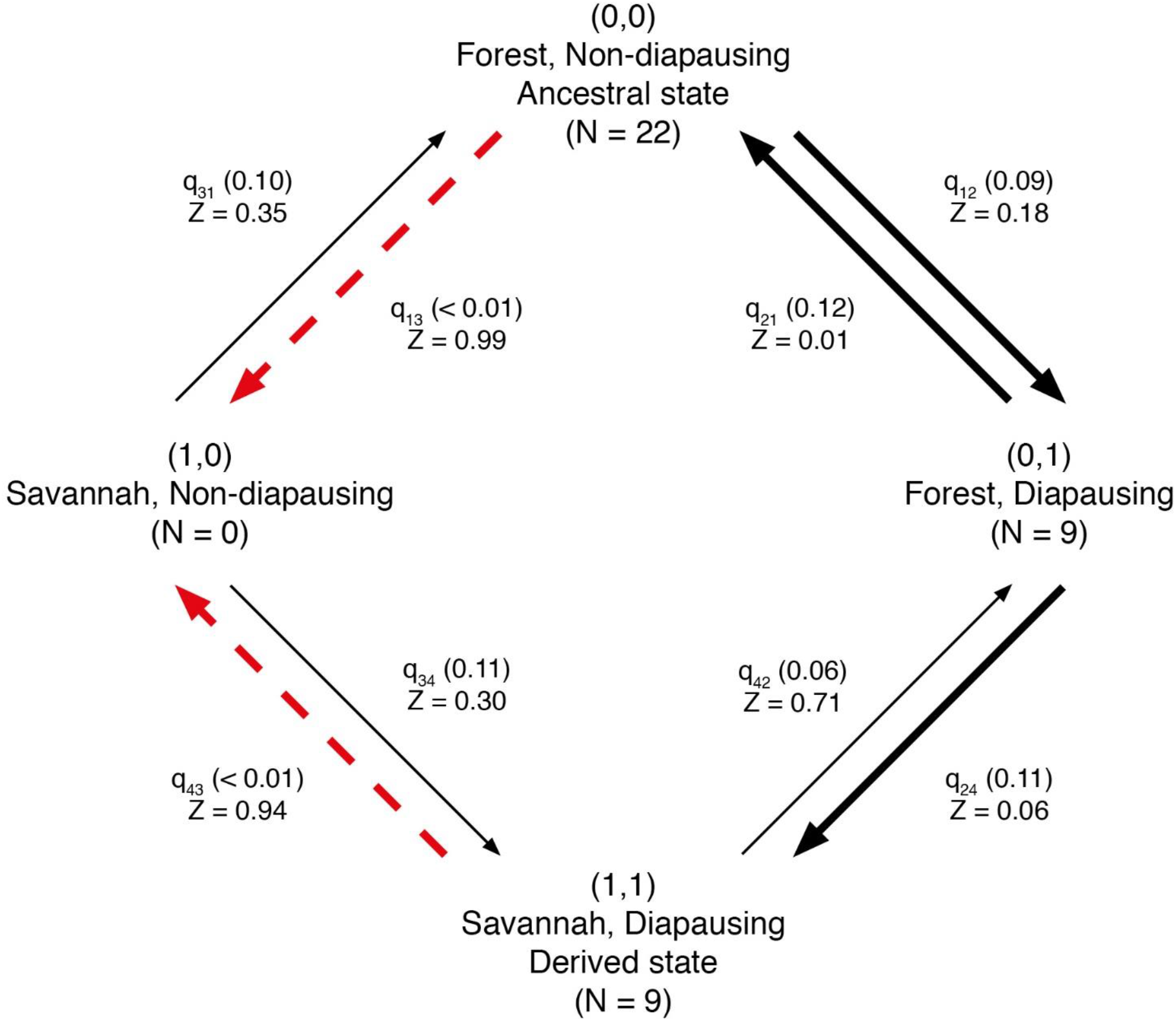
The evolutionary pathway depicting the habitat dependent evolution of the dry season mating strategies using 50% threshold and exponential hyperprior 0-10. The arrows represent three rate classes: thick black arrows are highly likely transitions between the states (Z value < 0.20); thin black arrows are the transitions with low confidence (Z value from 0.20-0.80); dashed red arrows represent unlikely transitions (Z value >0.80). The mean of posterior distribution of transition parameter are provided in the parentheses.

## 4. Discussion

Seasonal cycles in the tropics result in a highly uneven distribution of resources, including a minimal availability of resources in the dry season. Also, the extent of seasonality can vary across habitats, and forests may buffer extreme climatic variation across seasons while savannahs are highly seasonal (Archibald & Scholes, 2007). These types of dynamics can affect reproductive strategies in insects (Braby, 1995; Jones, 1987; Moore, 1986; Tauber et al., 1986). For example, populations or species inhabiting areas where larval hostplants are present throughout the year reproduce continuously, while they diapause where hostplant availability is seasonal (Braby, 1995; Dingle, 1968; Jones & Rienks, 1987). Such differential distribution of larval host plants in both habitats is likely to have led to the evolution of habitat-dependent mating strategies in Mycalesina butterflies. Adults of *Bicyclus* butterflies show three different dry season reproductive strategies: non-diapausing, diapausing, and pre-diapause mating. The incidence of these strategies varies across habitats. In general, the majority of forest-restricted species do not diapause, while all the savannah species show one of the two strategies involving some sort of reproductive diapause. Differences in strategies among species can be attributed to the phenology of host plants.

Some *Bicyclus* species of savannah habitats exhibit a pre-diapause mating strategy where females typically mate at the start of the dry season, store and perhaps utilise spermatophores, and then produce eggs at the onset of the rains. Two adaptive hypotheses have been proposed for this type of strategy. Firstly, spermatophores could provide nutrients to the female which may increase survival in the harsh dry season (Kato, 1986; Kato, 1989; Konagaya & Watanabe, 2015). This is especially relevant for savannah species as nutrients provided by spermatophores may favour female longevity until the end of the dry season (Svärd & Wiklund, 1988). Secondly, if the probability of encounters with potential mates after diapause is low due to high dry season mortality of males, mating before the onset of diapause would reduce the risk of being unfertilized when larval host plant conditions are favourable for oviposition. With respect to *Bicyclus*, this enables females to lay eggs immediately at the end of the dry season without any time lag in finding mates. Pre-diapause mating has been shown to occur in the Japanese population of a butterfly *Eurema mandarina* which shows a diapausing autumn-morph and a non-diapausing summer-morph (Kato, 1986; Kato, 1989; Konagaya & Watanabe, 2015). In this species, autumn-morph females mate with the summer-morph males at the start of the winter. In contrast, autumn-morph males diapause and show reproductive activity only at the end of the winters, and hence females may mate again with the same morph males. Whether this scenario is similar in *Bicyclus* butterflies is unclear, but it is possible that the dry season form females may mate with the wet season form males as there can be some overlap in flight between both forms. However, complete reproductive diapause could still be beneficial for females as mating at the end of the dry season means that their mate has expressed a strong dry season survival ability. Such a high mate quality might be more important than the potential benefits of the pre-diapause mating in some habitats, explaining why both strategies appear throughout *Bicyclus*.

In our long-term field samples, *B. safitza* exhibited an interesting pattern in which about one-quarter of the females had mature eggs in the peak dry season. The data show that the dry season form females tend to mate more often and have mature eggs in the early part, rather than in the middle of the dry season. This could be a bet-hedging strategy where a small proportion of females are reproductively active, while others undergo reproductive diapause (for example, Bradford & Roff, 1993). *B. safitza* predominantly prefers C_4_ grasses but in the dry season some individuals shift their preference towards utilization of C_3_ grasses in the forest (Nokelainen, Ripley, van Bergen, Osborne, & Brakefield, 2016; van Bergen et al., 2016). Hence, it is possible that a few reproductively active females found in the dry season might have shifted their preference to C_3_ grasses. Such mixed strategies can only be fully identified properly by longitudinal analysis of large number of samples in multiple dry seasons which is lacking in our museum samples. With few exceptions, a majority of our sampled species from the museum and smaller field collections showed a much clearer pattern suggesting that potential mixed strategies are rare. Two forest species, *B. dorothea* and *B. procora*, had 50% of spermatophores and eggs. There is evidence that time to reproductive maturity is longer in forest species (Braby, 2002) and hence detecting reproductive activity in 50% of specimens in these species may be due to delayed mating and slower maturation of eggs. On the contrary, if the species undergoes complete reproductive diapause, it is expected that all the females should be reproductively inactive. Our samples showed that a small percentage of females were mated with mature eggs but testing the robustness of our co-evolutionary models by setting narrow thresholds of 10% and 20% to classify species as diapausing consistently provided support for correlated evolution between habitat preference and mating strategies (supplementary material, Table S6).

Ancestral state reconstructions for habitat preference show that the ancestral *Bicyclus* were most likely forest-restricted species. This is in accordance with other studies demonstrating that forest habitats were ancestral followed by a more recent expansion of savannahs in the Late Miocene (Cerling et al., 1996; Edwards et al., 2010). In our analyses, the more ancestral clades are largely dominated by forest-restricted species, but there have been multiple independent colonization events into drier and more seasonal habitats during the Late Miocene and Pliocene. Similarly, for mating strategies, two diapause related strategies evolved during a similar time frame to that of habitat preferences. This congruent pattern suggests that the Late Miocene expansion of savannahs led to a rapid turnover in habitat preference and mating strategies in *Bicyclus* butterflies.

The evolution of key traits or innovations allows species to exploit novel niches and can also facilitate diversification (Hunter, 1998). It has been hypothesised that expansion of savannahs during the Late Miocene opened up new niches and their eventual colonization triggered diversification in the Mycalesina butterflies (Brakefield, 2010). Such savanna-linked diversification dynamics has been documented in several taxonomic groups (Aduse-Poku, Vingerhoedt, & Wahlberg, 2009; Cerling et al., 1997; De Silva, Peterson, Bates, Fernando, & Girard, 2017; Davis, Bakewell, Hill, Song, & Mayhew, 2018; Fuchs, Johnson, & Mindell, 2015; Kergoat et al., 2018; La Rosa, Pozio, & Hoberg, 2006; MacFadden & Hulbert, 1988; van Velzen, Wahlberg, Sosef, & Bakker, 2013; Zarlenga, Rosenthal, Toussaint et al., 2012). *Bicyclus* and other butterfly genera (e.g. *Cymothoe* and *Charaxes*, as well as the Satyrinae tribe in general) show a comparable burst of diversification during the Late Miocene which coincides with the expansion of savannahs (Aduse-Poku et al., 2009; Peña & Wahlberg, 2008; van Velzen et al., 2013). *Cymothoe* and *Charaxes* are generally forest species and diversification in this genus is hypothesised to be due to allopatric speciation driven by extensive forest fragmentation caused by savannah expansion (Aduse-Poku et al., 2009; van Velzen et al., 2013). However, in *Bicyclus* the forest species have radiated into savannahs and colonizing these habitats may have required novel adaptations, especially to survive in the resource-depleted dry season environment compared to the stable ancestral forest habitat. Though many studies have explored the link between the expansion of savannahs and diversification dynamics, potential key trait(s) that may have facilitated initial colonization of these habitats have seldom been explored. For example, the comparative studies of tooth morphology from fossil horses show that the grazing species from the Late Miocene had higher crown length than ancestral browsing species suggesting such adaptations may have helped in efficient grazing of tough C_4_ grasses (MacFadden, 2005; MacFadden & Hulbert, 1998). Similarly, grasshoppers feeding on C_3_ and C_4_ grasses show different mandibular morphology (Patterson, 1984), but this study lacked a phylogenetic approach. These studies reveal probable adaptations in herbivores linked to change in the vegetation structure during the Late Miocene in Africa.

The evolutionary pathway for *Bicyclus* species suggests that the gain of a capacity to undergo reproductive diapause in forest species preceded the habitat shift to savannahs (Figure 3). It is therefore possible that the initial gain of diapause in forest species evolved to utilise drier forest edges which likely became a more common habitat as a result of extensive fragmentation during the savannah expansions (Jacobs, 2004). However, ancestral state reconstructions suggest that around 10 million years ago the ancestors of the species that later colonized savannahs were generally forest-dependent species, but without a capacity to diapause (Figure 2). Taking this into consideration it appears that diapause was not strictly necessary to use forest edge habitats, but this trait evolved quickly multiple times when *Bicyclus* began to inhabit more open and seasonal habitats 5 to 10 million years ago. Given the multitude of current forest-dependent species that follow a diapausing strategy, it is likely that a continuing drying out of forest edge habitats played a key part in the evolution of reproductive diapause, and that having this ability further enabled *Bicyclus* species to eventually colonize novel savannah habitats.

## Acknowledgement

We would like to thank John Wilson for carrying out extensive sampling in Zomba, the samples which were used in the current study. Mark Williams and Ashanti African Tours arranged all logistics for the work in Ghana. Elishia Harji, Opuku Agyemang, Andrew Amankwaa and Osei ‘Yaw’ Johnson assisted with the field work in Ghana. The Wildlife division of Ghana Forestry Commission provided the necessary collection and export permits (0174076/024992) and the field work in Kenya was done in collaboration with the Insect Committee of Nature Kenya. This study was funded by ERC (Grant No. 250325) and John Templeton Foundation (Grant No. 60501) to PMB.

---

**Table S1:** Assigning mating strategies for museum data using 10%, 20%, and 50% threshold (ABRI-African Butterfly Research Institute; for habitat: FR, forest-restricted; FD- forest- dependent; O-open; for mating strategies: ND, non-diapausing; PDM, Pre-diapause mating; D, diapausing)

| Species | Habitat | Location | N | % mated | % eggs | 10% threshold | 20% threshold | 50% threshold |
| --- | --- | --- | --- | --- | --- | --- | --- | --- |
| *B taenias* | FD | ABRI | 6 | 0 | 0 | D | D | D |
| *B funebris* | FD | ABRI | 10 | 10 | 10 | D | D | D |
| *B italus* | FD | ABRI | 10 | 20 | 20 | ND | D | D |
| *B sausseri* | FD | ABRI | 10 | 70 | 70 | ND | ND | ND |
| *B anisops* | FD | ABRI | 10 | 90 | 90 | ND | ND | ND |
| *B sebetus* | FR | ABRI | 10 | 0 | 0 | D | D | D |
| *B procora* | FR | ABRI | 10 | 50 | 50 | ND | ND | ND |
| *B xeneas* | FR | ABRI | 10 | 63 | 63 | ND | ND | ND |
| *B auricruda* | FR | ABRI | 10 | 80 | 80 | ND | ND | ND |
| *B ignobilis* | FR | ABRI | 10 | 90 | 90 | ND | ND | ND |
| *B abnormis* | FR | ABRI | 9 | 100 | 100 | ND | ND | ND |
| *B dubia* | FR | ABRI | 9 | 100 | 100 | ND | ND | ND |
| *B evadne* | FR | ABRI | 10 | 100 | 100 | ND | ND | ND |
| *B maesseni* | FR | ABRI | 9 | 100 | 60 | ND | ND | ND |
| *B makomensis* | FR | ABRI | 9 | 100 | 100 | ND | ND | ND |
| *B uniformis* | FR | ABRI | 10 | 100 | 100 | ND | ND | ND |
| *B larseni* | FR | ABRI | 10 | * | 90.0 | ND | ND | ND |
| *B angulosa* | O | ABRI | 10 | 0 | 0 | D | D | D |
| *B cooksoni* | O | ABRI | 10 | 0 | 0 | D | D | D |
| *B cottrelli* | O | ABRI | 10 | 0 | 0 | D | D | D |
| *B pavonis* | O | ABRI | 10 | 60 | 20 | ND | PDM | PDM |
| *B milyas* | O | ABRI | 10 | 67 | 10 | PDM | PDM | PDM |
| *B anynana* | O | ABRI | 19 | 89 | 11 | PDM | PDM | PDM |
| *B mandanes* | FD | Field (Ghana) | 12 | 0 | 0 | D | D | D |
| *B vulgaris* | FD | Field (Ghana) | 61 | 0 | 0 | D | D | D |
| *B funebris* | FD | Field (Ghana) | 91 | 1 | 0 | D | D | D |
| *B zinebi* | FD | Field (Ghana) | 8 | 38 | 38 | ND | ND | D |
| *B dorothea* | FD | Field (Ghana) | 20 | 50 | 50 | ND | ND | ND |
| *B martius* | FD | Field (Ghana) | 21 | 57 | 50 | ND | ND | ND |
| *B sandace* | FD | Field (Ghana) | 42 | 64 | 64 | ND | ND | ND |
| *B madetes* | FD | Field (Ghana) | 34 | 74 | 14 | ND | PDM | PDM |
| *B sambulos* | FR | Field (Ghana) | 10 | 30 | 10 | PDM | PDM | D |
| *B xeneas* | FR | Field (Ghana) | 11 | 91 | 91 | ND | ND | ND |
| *B safitza* | O | Field (Ghana) | 24 | 8 | 4 | D | D | D |
| *B smithi* | FD | Field (Kenya) | 37 | 89 | 89 | ND | ND | ND |
| *B jeffryi* | FD | Field (Kenya) | 10 | 90 | 90 | ND | ND | ND |
| *B buea* | FD | Field (Kenya) | 12 | 100 | 92 | ND | ND | ND |
| *B golo* | FR | Field (Kenya) | 7 | 86 | 86 | ND | ND | ND |
| *B mollitia* | FR | Field (Kenya) | 7 | 100 | 100 | ND | ND | ND |
| *B campa* | O | Field (Nigeria) | 11 | 27 | 0 | PDM | PDM | D |
| *B campina* | FD | Field (Zomba) | 232 | 65 | 65 | ND | ND | ND |
| *B ena* | O | Field (Zomba) | 206 | 84 | 24 | ND | ND | PDM |
| *B safitza* | O | Field (Zomba) | 373 | 22 | 22 | ND | ND | D |

\* In *B. larseni* it was not possible to find spermatophores in most of the samples due to its small size (see main text more information)

**Table S2:** AIC weights for different models for ancestral state reconstruction of mating strategies and habitat preference (values in bold represent models with the highest AIC weights that were used for producing Figure 2 in the main text).

| States | Mating strategy | Habitat preference |
| --- | --- | --- |
| Equal rates model | **0.62** | **0.60** |
| Symmetric model | 0.34 | 0.33 |
| All rates different model | 0.03 | 0.07 |

**Table S3:** Marginal likelihood value and Bayes Factor for the independent and dependent models using three different priors (exponential hyperior 0-10 & 0-100; uniform 0-100) for assessing correlated evolution for three independent runs.

| **Exponential hyperprior 0 10** | **Independent** | **Dependent** | **Bayes Factor** |
| --- | --- | --- | --- |
| Run 1 | -47.48 | -40.54 | 13.88 |
| Run 2 | -47.45 | -40.48 | 13.94 |
| Run 3 | -47.54 | -40.43 | 14.22 |
| *Average* | *-47.49* | *-40.48* | *14.01* |
| **Uniform 0 100** | **Independent** | **Dependent** | **Bayes Factor** |
| Run 1 | -50.67 | -45.56 | 10.22 |
| Run 2 | -50.67 | -45.57 | 10.2 |
| Run 3 | -50.68 | -45.56 | 10.24 |
| *Average* | *-50.67* | *-45.56* | *10.22* |
| **Exponential hyperprior 0 100** | **Independent** | **Dependent** | **Bayes Factor** |
| Run 1 | -49.18 | -42.04 | 14.28 |
| Run 2 | -49.24 | -42.03 | 14.42 |
| Run 3 | -49.31 | -42.03 | 14.56 |
| *Average* | *-49.24* | *-42.03* | *14.42* |

**Table S4:** Marginal likelihood value and Bayes Factor for the independent and dependent models to test the effect of merging forest-dependent (FD) state with forest-restricted (FR) and open (O) habitat using exponential hyperprior 0-10.

|  | **FD merged to FR (original data set)** |  |  |  | **FD merged to O** |  |  |
| --- | --- | --- | --- | --- | --- | --- | --- |
|  | Independent | Dependent | Bayes Factor |  | Independent | Dependent | Bayes Factor |
| Run1 | -47.48 | -40.54 | *13.88* |  | -51.4 | -48.7 | *5.4* |
| Run2 | -47.45 | -40.48 | *13.94* |  | -51.68 | -48.73 | *5.9* |
| Run3 | -47.54 | -40.43 | *14.22* |  | -51.39 | -48.76 | *5.26* |
| *Average* | *-47.49* | *-40.48* | *14.01* |  | *-51.49* | *-48.73* | *5.52* |

**Table S5:**
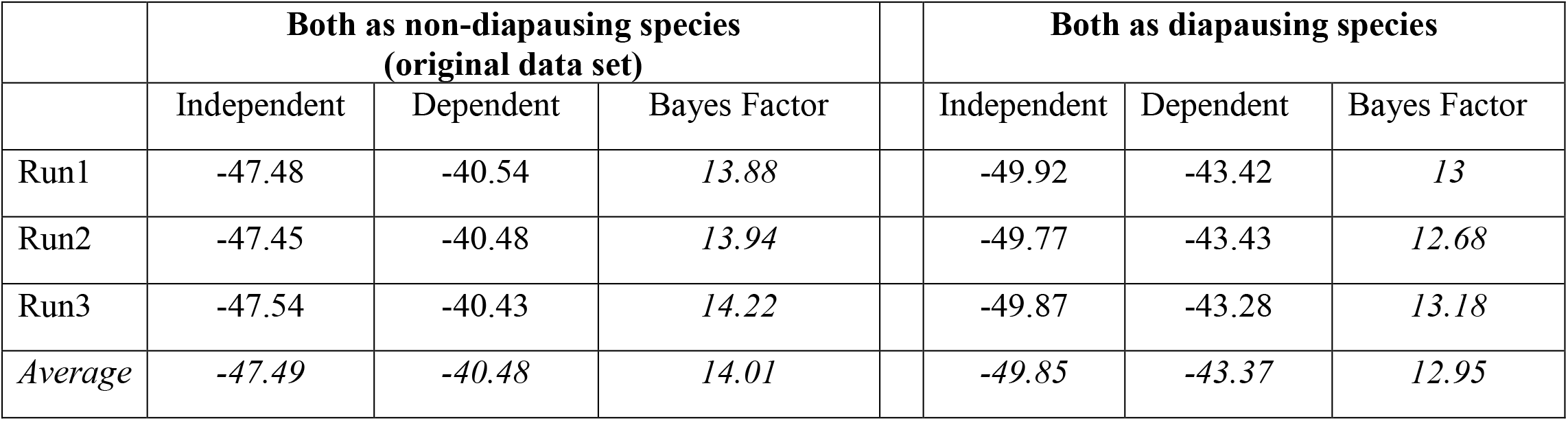
Marginal likelihood value and Bayes Factor for the independent and dependent models to test the effect of classifying two species *B. dorothea* and *B. procora* as non-diapausing and diapasing species using exponential hyperprior 0-10.

|  | **Both as non-diapausing species**<br>**(original data set)** |  |  |  | **Both as diapausing species** |  |  |
| --- | --- | --- | --- | --- | --- | --- | --- |
|  | Independent | Dependent | Bayes Factor |  | Independent | Dependent | Bayes Factor |
| Run1 | -47.48 | -40.54 | 13.88 |  | -49.92 | -43.42 | 13 |
| Run2 | -47.45 | -40.48 | 13.94 |  | -49.77 | -43.43 | 12.68 |
| Run3 | -47.54 | -40.43 | 14.22 |  | -49.87 | -43.28 | 13.18 |
| Average | -47.49 | -40.48 | 14.01 |  | -49.85 | -43.37 | 12.95 |

**Table S6:** Marginal likelihood value and Bayes Factor for the independent and dependent models to test the effect of two different thresholds on the classification of mating strategies using exponential hyperprior 0-10.

|  | **50%**<br>**(original data set)** |  |  |  | **10%** |  |  |  | **20%** |  |  |
| --- | --- | --- | --- | --- | --- | --- | --- | --- | --- | --- | --- |
|  | Independent | Dependent | Bayes Factor |  | Independent | Dependent | Bayes Factor |  | Independent | Dependent | Bayes Factor |
| Run1 | -47.48 | -40.54 | 13.88 |  | -45.9 | -43.36 | 5.08 |  | -49.2 | -46.12 | 6.16 |
| Run2 | -47.45 | -40.48 | 13.94 |  | -45.92 | -43.33 | 5.18 |  | -49.19 | -46.17 | 6.04 |
| Run3 | -47.54 | -40.43 | 14.22 |  | -45.93 | -43.32 | 5.22 |  | -49.19 | -46.19 | 6 |
| Avg. | -47.49 | -40.48 | 14.01 |  | -45.91 | -43.33 | 5.16 |  | -49.19 | -46.16 | 6.06 |

**Table S7:** Marginal likelihood value and Bayes Factor for the independent and dependent models to test the effect of sample size on correlated evolution using exponential hyperprior 0-10. For each set 30% of the species were randomly removed from the data set.

|  | Run1 |  |  |  | Run2 |  |  |  | Run3 |  |  |
| --- | --- | --- | --- | --- | --- | --- | --- | --- | --- | --- | --- |
|  | Independent | Dependent | Bayes Factor |  | Independent | Dependent | Bayes Factor |  | Independent | Dependent | Bayes Factor |
| Set1 | -36.41 | -32.38 | 8.06 |  | -36.4 | -32.31 | 8.18 |  | -36.38 | -32.32 | 8.12 |
| Set2 | -35.17 | -30.58 | 9.18 |  | -35.17 | -30.7 | 8.94 |  | -35.15 | -30.57 | 9.16 |
| Set3 | -36.79 | -31.75 | 10.08 |  | -36.73 | -31.75 | 9.96 |  | -36.82 | -31.77 | 10.1 |
| Set4 | -37.09 | -32.76 | 8.66 |  | -37.06 | -32.72 | 8.68 |  | -37.05 | -32.76 | 8.58 |
| Set5 | -36.86 | -31.65 | 10.42 |  | -36.84 | -31.62 | 10.44 |  | -36.86 | -31.64 | 10.44 |
| Set6 | -36.07 | -32.03 | 8.08 |  | -36.04 | -32 | 8.08 |  | -36.03 | -32.02 | 8.02 |
| Set7 | -36.69 | -30.86 | 11.66 |  | -36.73 | -30.85 | 11.76 |  | -36.72 | -30.81 | 11.82 |
| Set8 | -36.39 | -30.96 | 10.86 |  | -36.43 | -31 | 10.86 |  | -36.33 | -30.93 | 10.8 |
| Set9 | -36.07 | -31.33 | 9.48 |  | -36.04 | -31.28 | 9.52 |  | -36.14 | -31.26 | 9.76 |
| Set10 | -35.96 | -31.26 | 9.4 |  | -35.92 | -31.33 | 9.18 |  | -35.9 | -31.3 | 9.2 |
| Avg. | -36.35 | -31.55 | 9.85 |  | -36.33 | -31.55 | 9.56 |  | -36.33 | -31.53 | 9.6 |

**Figure S1:**
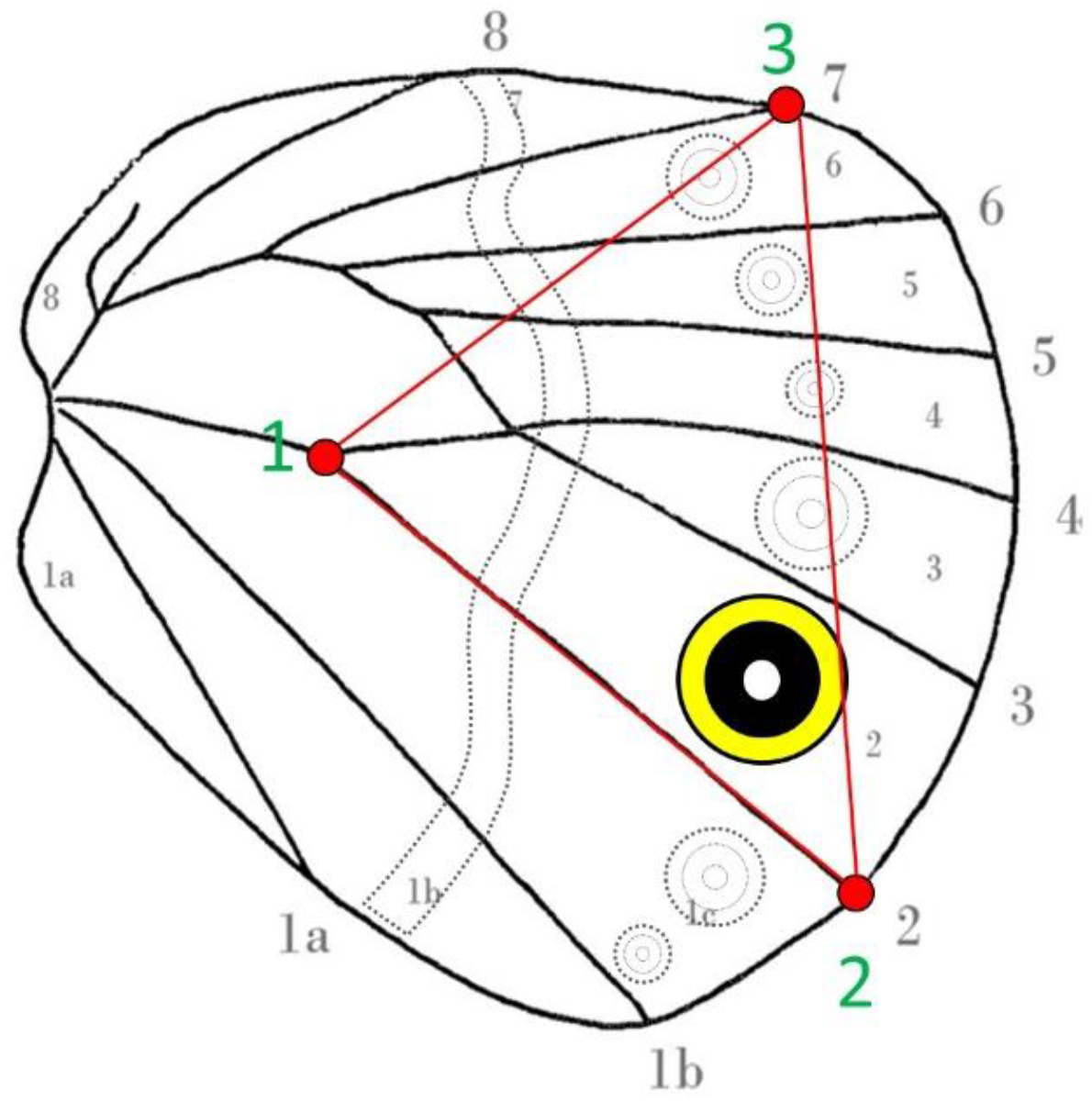
Traits measured on the ventral hind wing for classification of the wet and dry season forms. The triangular area within the three landmarks was used as a proxy for hindwing area and was used to calculate the relative area of the combined yellow, black and white regions of the CuA1 eyespot that is highlighted in colour.

**Figure S2:**
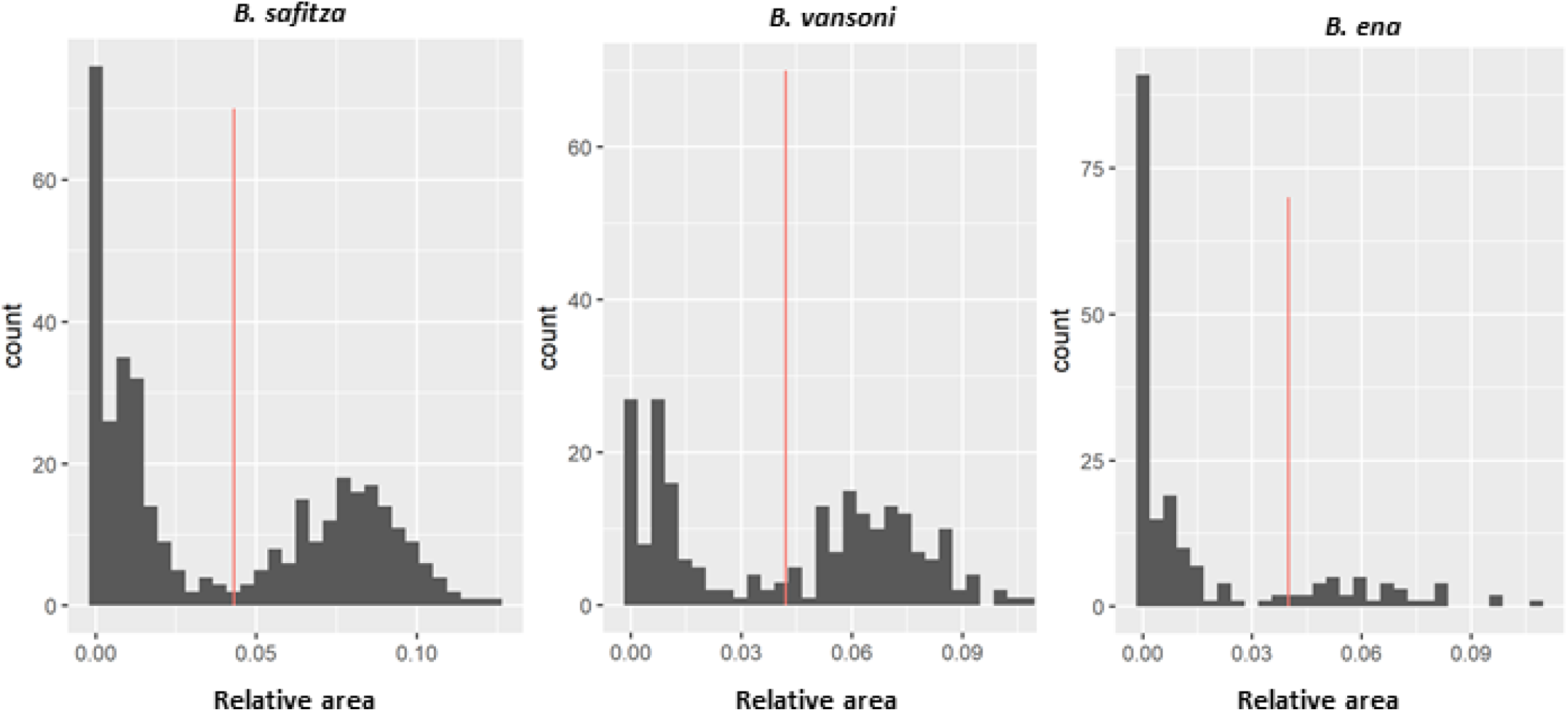
Classification of seasonal forms based on relative area of the ventral eyespot by setting a threshold value (red line); values above and below this threshold were classified as wet and dry season forms, respectively.

